# Alcohol disrupts long-term potentiation at hippocampus-medium spiny neuron synapses in the medial shell of the nucleus accumbens

**DOI:** 10.1101/2025.07.09.663974

**Authors:** Ashley E. Copenhaver, Sarah A. Snider, Jalane R. Campbell, Tara A. LeGates

## Abstract

Chronic alcohol exposure is a major driver of alcohol use disorders (AUD), in part through its ability to induce maladaptive plasticity within neural circuits that regulate reward, motivation, and affect. Excitatory projections from the hippocampus (Hipp) to the nucleus accumbens (NAc) play a pivotal role in regulating reward-related behaviors, and this pathway serves as a key locus for establishing associations between rewarding stimuli and related contextual information. Regulation of the strength of Hipp-NAc synapses is critical for supporting these behaviors, and impairments in Hipp-NAc plasticity are associated with anhedonia and disrupted reward learning. Here, we demonstrate that acute ethanol application to *ex vivo* brain slices prevents long-term potentiation (LTP) at Hipp-NAc synapses without altering presynaptic release probability. Furthermore, chronic intermittent exposure to ethanol abolishes LTP at these synapses, an effect we observed during early abstinence indicating that ethanol-induced disruptions in Hipp-NAc plasticity outlasts exposure periods. Together, our findings demonstrate that ethanol rapidly alters Hipp-NAc plasticity, and its effects are still evident in early abstinence. Given the behavioral relevance of these synapses, this work has important implications for the mechanisms underlying ethanol-dependent effects on reward processing and negative affective states associated with AUD.

## 1. INTRODUCTION

Alcohol exerts both immediate and long-lasting effects on motivated behaviors, cognitive function, and emotion regulation^1^. These effects arise primarily from alcohol’s direct modulation of synaptic transmission and its influence over plasticity within key brain reward circuits ^2^. Although heightened plasticity in reward pathways may initially promote alcohol-seeking behaviors, persistent changes in circuit activation can ultimately impair the brain’s ability to respond to natural rewards and trigger prolonged activation of stress-related neural systems. As a result, disruption of normal reward processing and persistent stress signaling are thought to underlie the negative emotional states frequently associated with chronic alcohol use and dependence. AUD with comorbid anhedonia (loss of pleasure derived from previously pleasurable activities) is associated with increased risk of relapse and poorer clinical outcomes, presenting a major obstacle to recovery^3,4^.

The NAc is a key mediator of motivated behaviors where both stress and rewarding experiences modulate synaptic function and plasticity to drive alterations in behavior^5–7^. Excitatory input from several key brain regions (e.g., hippocampus, prefrontal cortex, thalamus, amygdala) has a crucial role in regulating NAc neuron activity to effectively encode reward-associated information and modulate motivated behaviors^8–10^. Alterations in the strength and plasticity of these synapses as well as changes in NAc neuron excitability have been associated with substance use disorders and depression, further demonstrating the key role of NAc activity in modulating motivated behaviors and emotional states^11–13^. Chronic ethanol exposure and chronic stress produce convergent effects in the NAc, both altering excitability and disrupting plasticity, thereby promoting anhedonia and dysregulated reward processing^14–16^.

Projections from the hippocampus (Hipp) provide extensive excitatory input that has a major influence on NAc activity^10,17–19^. Hipp-NAc connectivity has been tied to several behaviors related to reward and motivation in humans and rodents including reinforcement, positive affect, as well as spatial- and contextual-based reward learning^20–31^. Our previous work revealed that activity-dependent strengthening of Hipp-NAc synapses drives reward-related behaviors while chronic stress disrupts this plasticity and induces anhedonia^24^. Hipp and NAc activity also influences alcohol-related behaviors, where reduced Hipp activity increases alcohol consumption, while activation of Hipp-NAc synapses attenuates excessive drinking^32^. Given that both stress and alcohol have been linked to reward dysregulation, we sought to determine whether ethanol also alters LTP at Hipp-NAc synapses. Focusing on medium spiny neurons (MSNs), which represent ∼95% of the neuron population in the NAc, we found that both acute *ex vivo* and chronic *in vivo* exposure to ethanol prevents LTP at Hipp-NAc synapses in males and females. Given the link between Hipp-NAc potentiation in mediating reward-related behaviors, ethanol’s effects on these synapses may be a key mechanism driving maladaptive reward-related behaviors observed in response to alcohol use.

## 2. MATERIALS AND METHODS

Male and female C57BL/6J mice (8-10 weeks) were group housed (2-5 mice per cage) in a 12:12LD cycle with food and water ad libitum. We did not track the estrous cycle in female mice. All experiments were performed in accordance with the regulations set forth by the Institutional Animal Care and Use Committee at the University of Maryland, Baltimore County.

### 2.1. Ethanol Vapor Chamber Exposure

Male and female mice (8-10 weeks at the start of the experiment) were placed in acrylic inhalation chambers (60 × 59 × 35 cm) in their home cages and exposed to chronic intermittent ethanol (CIE) vapor or air 14 hours/day (ZT11-1), 4 days a week, for a total of 3 weeks. Compressed air (10–20 psi) was passed through a tube submerged in 95% ethanol to volatilize. Control vapor chambers delivered only air with no ethanol vapor. Vapor chamber ethanol concentrations were monitored daily, and air flow was adjusted to produce ethanol concentrations within 1.8-2.0% ethanol content as measured by a digital alcohol breath tester (BACTrack). After four days in the inhalation chambers, mice underwent a 72-h forced abstinence period. Whole cell electrophysiology was performed on week 4 after the 72-h abstinence period (3-6 days following last ethanol exposure). Mice did not receive pyrazole injections to avoid potential stress-related confounds associated with pyrazole administration^33,34^. Pyrazole-free paradigms has been previously described as a valid alternative to pyrazole-assisted protocols^33–35^.

### 2.2. Blood ethanol concentration (BEC) measurements

Immediately following the last day of exposure each week, mice were removed from the vapor chamber, and blood was sampled via a tail nick. Blood was collected in capillary tubes and then transferred to 1.5mL tubes. Samples were centrifuged, and serum was transferred to new tubes. BEC was measured with an AM1 Alcohol Analyzer.

### 2.3. Brain Slice Preparation

Acute parasagittal slices (lateral 0.36-0.72) containing the fornix and nucleus accumbens were prepared for whole-cell patch-clamp electrophysiology. Animals were deeply anesthetized with isoflurane, decapitated, and brains were quickly dissected and submerged in ice-cold, bubbled (carbogen: 95% O_2_/5% CO_2_) N-methyl-D-glucamine (NMDG) recovery solution containing the following (in mM): 93 NMDG, 2.5 KCl, 1.2 NaH_2_PO_4_, 11 glucose, 25 NaHCO_3_, 1.2 MgCl_2_, and 2.4 CaCl_2_, pH=7.3-7.4, osmolarity=300-310 mOsm. Using a vibratome (VT1000S, Leica Microsystems), parasagittal slices (400 μm) were cut in cold, oxygenated NMDG. Slices were transferred to 32-34°C NMDG for 7-12 minutes to recover and were then transferred to room-temperature artificial cerebrospinal fluid (aCSF) containing the following (in mM): 120 NaCl, 3 KCl, 1.0 NaH_2_PO_4_, 20 glucose, 25 NaHCO_3_, 1.5 MgCl_2_·7H_2_O, and 2.5 CaCl_2_, pH=7.3-7.4. Slices were allowed to recover for 1-hour at room-temperature before beginning electrophysiological recordings.

### 2.4. Whole-Cell Recordings

Slices were placed in a submersion-type recording chamber and superfused with room-temperature aCSF (flow rate 0.5-1mL/min). Cells were visualized using a 60x water immersion objective on a Nikon Eclipse FN-1 or Scientifica microscope. Whole-cell patch-clamp recordings were performed using an Axopatch 200B amplifier (Axon Instruments, Molecular Devices) and a Digidata 1550B digitizer (Axon Instruments). Patch pipettes (4-8MΩ) were made from borosilicate glass (World Precision Instruments) using a Sutter Instruments P-97 model puller.

All recordings were performed in voltage-clamp from MSNs in the NAc medial shell. A bipolar stimulating electrode (FHC) was placed in the fornix to electrically stimulate hippocampal axons while recording evoked excitatory postsynaptic currents (EPSC) (Figure 1a). Patch pipettes were filled with a solution containing 130 mM K-gluconate, 5 mM KCl, 2 mM MgCl6-H2O, 10 mM HEPES, 4 mM Mg-ATP, 0.3 mM Na2-GTP, 10 mM Na2-phosphocreatine, and 1 mM EGTA; pH=7.3-7.4; osmolarity=285-295mOsm. EPSCs were recorded at -70mV from paired pulses (100ms apart) elicited every 10s. After obtaining a stable 5-minute baseline EPSC recording, high-frequency stimulation (HFS: four trains of 100Hz stimulation for 1s with 15s between trains while holding the cell at -40mV) was applied, followed by a 30-minute recording of EPSCs. One cell was recorded per slice. For acute ethanol experiments, aCSF containing 50mM ethanol (Sigma) was washed onto slices for at least 20 minutes prior to recording, and experiments were done in the presence of ethanol. We used the following exclusion criteria to eliminate unhealthy cells and unreliable recordings: 1) We only proceeded with experiments on cells with series resistances <10MΩ, 2) Cells were excluded if series resistance changed by >20% (comparing the resistance at the beginning and end of the experiment), 3) Cells in poor health or poor recording status were excluded (i.e. cell partially or fully sealed up, a decrease in holding current >100pA that is consistent with the cell dying, an increase in the variability of timing of EPSCs (jitter) post HFS, and/or an increase in response failure rate to > 50%).

**Figure 1:**
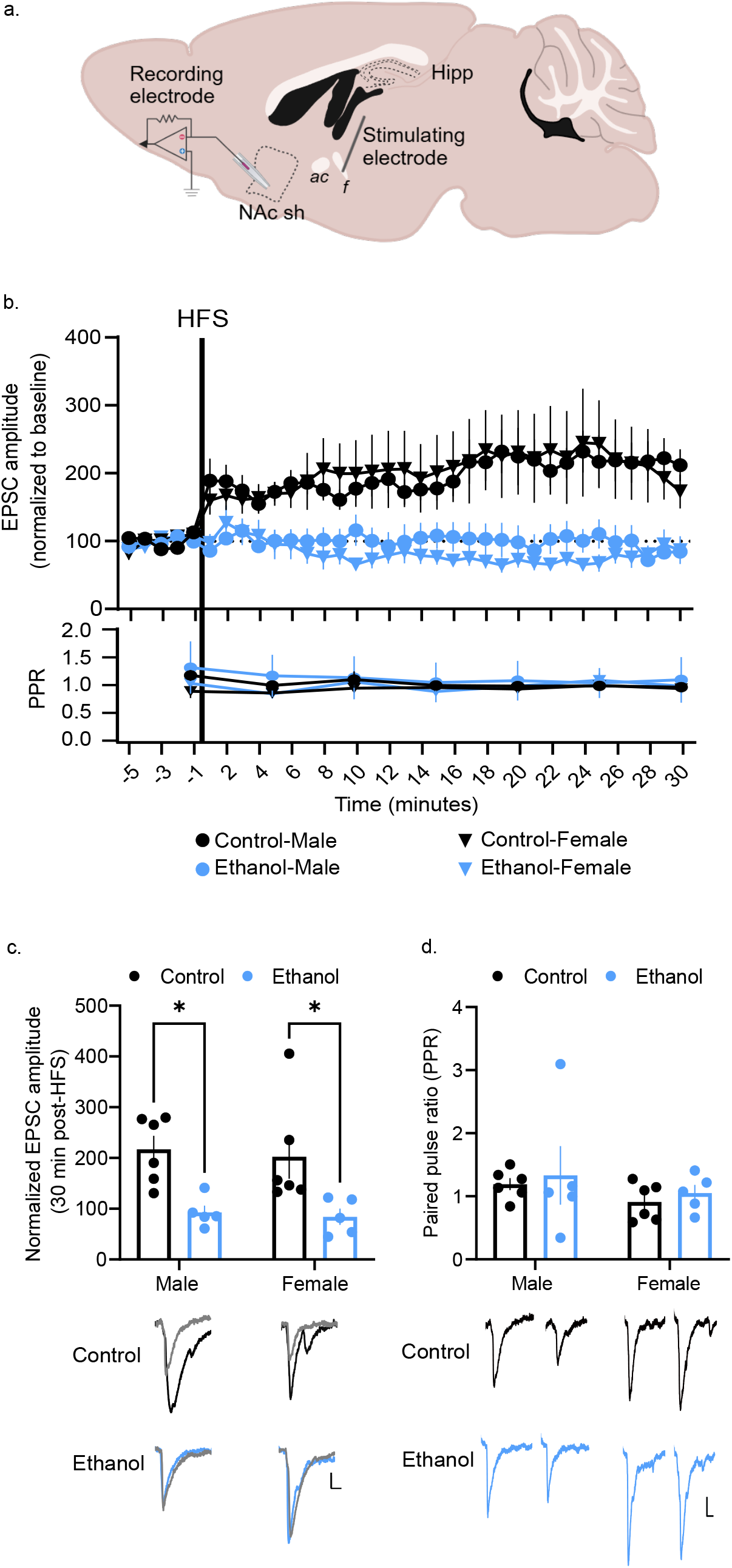
Ethanol interferes with LTP at Hipp-NAc synapses. a) Schematic of recording strategy with stimulating electrode in the fornix and recording electrode in the NAc medial shell to record Hipp-evoked EPSCs from MSNs. The illustrated parasagittal section represents a slice from lateral +0.48mm. ac-anterior commissure f-fornix. b) Pretreatment of slices with 50mM ethanol prevents HFS-induced LTP at Hipp-NAc synapses in males and females (top). Paired-pulse ratios were not altered by HFS or ethanol (bottom). c) Summary data from the last 5 minutes of recording with representative traces below. Gray traces indicate baseline and black/blue indicate post-HFS. d) No differences in paired-pulse ratios recorded in the presence and absence of ethanol with representative traces shown below. n= 6_control male_, 4_control female_, 5_ethanol male_, 5_ethanol female_ cells. * p<0.05, Scale bar = 20pA/10ms.

### 2.5. Data analysis

To analyze data from HFS experiments, EPSC amplitudes were normalized to the average baseline amplitude. Normalized EPSC amplitudes were averaged over the last 5 minutes of the recording (minutes 25-30 post HFS) to examine the direction and magnitude of plasticity. Paired pulse ratio (PPR) was measured by calculating 5-minute averages of EPSC amplitudes from our paired pulses (EPSC_1_ and EPSC_2_) and dividing EPSC_2_ by EPSC_1_ (ie: EPSC_2_/EPSC_1_ = PPR). Statistical tests were performed using GraphPad Prism 9/10 software. A p-value of <0.05 was considered statistically significant, and exact p-values can be found in the results section. For acute ethanol experiments, n represents the number of cells, which is equivalent to the number of slices as one LTP experiment is performed per slice. For CIE experiments, n represents the number of mice. For line graphs, points represent the mean and error bars indicate SEM. Summary data plots display one point for each cell, with each point representing the average EPSC amplitude from the last 5 minutes of recording. For bar graphs, the bar represents the mean and error bars indicate SEM with individual data points overlayed. Statistical significance (p<0.05) was assessed by Two-way ANOVA (sex x treatment) with Šídák’s multiple comparisons. One-sample Wilcoxon tests were used to determine whether average EPSC amplitude from the last 5 minutes of recording was significantly different than baseline, indicating LTP. A three-way ANOVA (sex x treatment x time) was used to analyze PPR data before and after HFS. Two-way RM ANOVA was used to compare BECs from males and females over the course of CIE.

## 3. RESULTS

### 3.1. Acute ethanol application to *ex vivo* slices prevents Hipp-NAc LTP

To determine whether ethanol acutely influences plasticity at Hipp-NAc synapses, we recorded HFS-induced LTP in the presence and absence of 50mM ethanol (Figure 1a). This concentration was chosen because it reliably interferes with NMDAR and VGCC function^36,37^, which we previously demonstrated are required for Hipp-NAc LTP in males and females, respectively^38^. We found that, in the absence of ethanol, HFS induced LTP at Hipp-NAc synapses in slices taken from male and female mice with no difference in magnitude between the sexes (Figure 1b-c), similar to our previous published results^24,38^. However, pretreatment with 50mM ethanol prevented LTP induction in slices taken from males and females (Figure 1b-c; Sex x Treatment Interaction: F_(1,18)_ = 0.01, p=0.92; Main effect of Sex: F_(1,18)_ = 0.15, p=0.70; Main effect of treatment: F_(1,18)_ = 16.65, p=0.0007; Šídák’s multiple comparisons post hoc: p_(Male:control vs EtOH)_ = 0.017, p_(Female:control vs EtOH)_ = 0.02; One-sample Wilcoxon test: Theoretical median=100, Control Male: W=21, p=0.03, Control Female: W=21, p=0.03, Ethanol Male: W=-5, p=0.63, Ethanol Female: W=-5, p=0.63; n= 6_control male_, 6_control female_, 5_ethanol male_, 5_ethanol female_ cells.). We observed no difference in baseline paired pulse ratio (PPR) (Figure 1d; Sex x Treatment Interaction: F_(1,18)_ = 7.86e-006, p=0.99; Main effect of Sex: F_(1,18)_ = 1.47, p=0.24; Main effect of treatment: F_(1,18)_ = 0.36, p=0.56) suggesting that application of ethanol did not alter basal release probability. Furthermore, no change in PPR was observed following HFS irrespective of whether ethanol was present (Figure 1b, bottom; Time x Sex x Treatment Interaction: F_(2.621, 47.18)_ = 0.44, p=0.70; Time x Treatment Interaction: F_(2.621, 47.18)_ = 2.19, p=0.11; Time x Sex Interaction: F_(2.590, 41.45)_ = 0.38, p=0.74), suggesting that HFS did not elicit changes in presynaptic neurotransmitter release probability.

### 3.2. CIE disrupts Hipp-NAc LTP

To determine whether chronic exposure to ethanol leads to changes in Hipp-NAc plasticity, we used CIE exposure to ethanol vapor for controlled delivery and dosing of ethanol to mice over a 3-week period (Figure 2a). Because previous work has shown that stress negatively impacts plasticity of these synapses^24,39,40^, CIE was done without pyrazole injections to order to isolate the effects of ethanol exposure itself and minimize confounding effects of repeated injections and pyrazole-related stress responses^33,34^. BECs measured weekly were within the low range previously associated with ethanol-induced changes in reward-related behaviors^41,42^. BECs appeared to decrease over time, but this was only statistically significant in females (Figure 2b; Sex x Time Interaction: F_(2,18)_=0.69, p=0.52; Main effect of sex: F_(1,9)_=0.13, p=0.73; Main effect of time: F_(2,18)_=8.61, p=0.002; n=5 males and 6 females). Following at least 72 hours of forced abstinence, slices from CIE-exposed mice demonstrated a deficit in HFS-induced LTP relative to air-exposed controls, which showed typical LTP (Figure 2c-d; Sex x Treatment Interaction: F_(1,20)_ = 0.04, p=0.85; Main effect of Sex: F_(1,20)_ = 0.11, p=0.74; Main effect of treatment: F_(1,20)_ = 12.53, p=0.002; Šídák’s multiple comparisons post hoc: p_(Male:control vs EtOH)_ = 0.04, p_(Female:control vs EtOH)_ = 0.04; One-sample Wilcoxon test: Theoretical median=100, Control Male: W=28, p=0.02, Control Female: W=21, p=0.03, Ethanol Male: W=-1, p>.99, Ethanol Female: W=-1, p>.99; n= 7_control male_, 6_control female_, 6_ethanol male_, 5_ethanol female_ mice). This effect was consistent in both male and female mice. Comparison of baseline PPRs revealed no difference between CIE mice and air exposed controls (Figure 2e; Sex x Treatment Interaction: F_(1,20)_ = 0.23, p=0.64; Main effect of Sex: F_(1,20)_ = 0.0009, p=0.98; Main effect of treatment: F_(1,20)_ = 1.15, p=0.30), suggesting that CIE did not induce changes in release probability. Similar to the data shown in Figure 1, HFS did not induce changes in PPR regardless of ethanol exposure or sex (Figure 2c bottom; Time x Sex x Treatment Interaction: F_(6,20)_ = 1.60, p=0.15; Time x Treatment Interaction: F_(6,20)_ = 0.65, p=0.69; Time x Sex Interaction: F_(6,20)_ = 1.07, p=0.39), demonstrating a lack of effect on presynaptic function.

**Figure 2:**
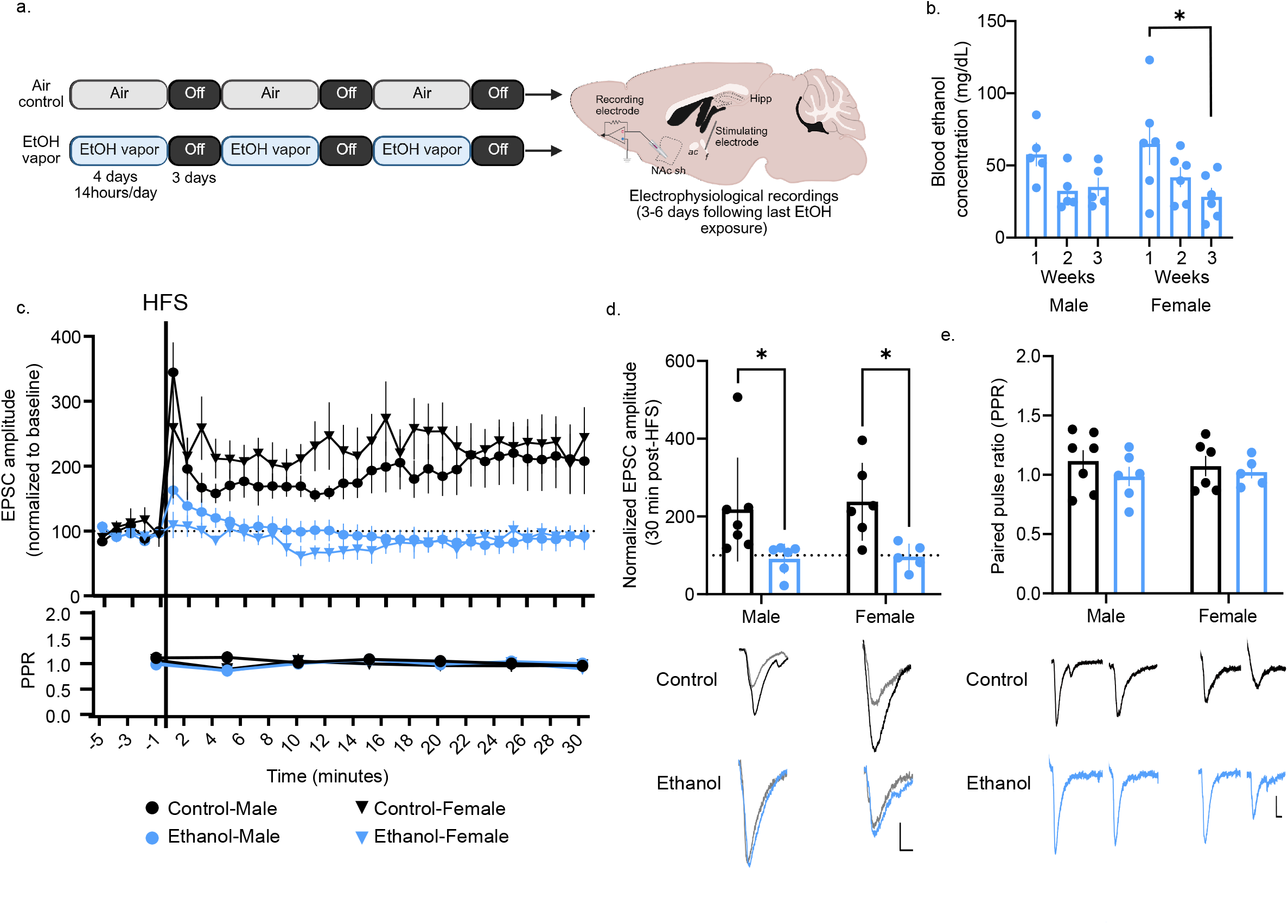
Chronic intermittent ethanol exposure disrupts LTP at Hipp-NAc synapses. a) Schematic of experimental design to record Hipp-evoked EPSCs in the NAc shell of mice exposed to CIE or air. ac-anterior commissure f-fornix. b) BECs from male and female mice exposed to CIE. (n=5 male, 6 female.) c) Slices taken from male and female mice exposed to CIE show disrupted LTP as compared to air-exposed controls (top). Paired-pulse ratios were not altered by HFS or ethanol (bottom). d) Summary data from the last 5 minutes of recording with representative traces below. Gray traces indicate baseline and black/blue indicate post-HFS. e) No differences in paired-pulse ratios between cells recorded from air- and CIE-exposed mice with representative traces shown below. n= 7_control male_, 6_control female_, 6_ethanol male_, 5_ethanol female_ mice * p<0.05, Scale bar = 20pA/10ms.

## 4. DISCUSSION

Acute alcohol intoxication and chronic alcohol consumption have extensive effects on cognitive processing, motivation, and affective state. Bidirectional regulation of the strength of Hipp-NAc connectivity plays a central role in regulating reward-related learning and affect and has been implicated in substance use and depression^23–26,28,29,32^. We previously demonstrated that LTP of Hipp-NAc synapses facilitates reward-related behaviors while disruption is associated with anhedonia^24^. Here, we show that both acute and chronic ethanol exposure also interfere with LTP at these synapses, presenting a potential site through which alcohol alters motivation and affect.

Alcohol’s ability to influence cognitive and emotional function across timescales is likely driven, in part, by its effects on synaptic plasticity within key circuits^43–46^. Our data demonstrate that acute application of ethanol to brain slices and chronic *in vivo* exposure disrupt LTP at Hipp-NAc synapses similarly in males and females. We previously demonstrated LTP at these synapses is mediated by sex-specific mechanisms: LTP is NMDAR-dependent in males and L-type VGCCs-dependent in females^38^. Given that ethanol can interfere with NMDAR- and VGCC-dependent signaling^14,36,43,47^, dual actions on NMDAR- and VGCC-mediated plasticity offer one potential explanation for LTP impairments observed across sexes.

Recent work has shown that ethanol disrupts NMDAR-dependent LTD in the NAc, an effect specific to D1-MSNs^14,15,48^. We did not identify MSN subtypes, which is a limitation given these subtype-specific effects, and future work on identified subtypes will be necessary for a full mechanistic understanding of ethanol’s effects. Ethanol-induced disruption of LTD at Hipp-NAc synapses coincided with both presynaptic and postsynaptic alterations^14^, implicating both pre- and postsynaptic mechanisms in ethanol-induced plasticity deficits. In contrast, we observed no differences in PPR no detectable change in presynaptic release (Figure 2e). These differences in experimental outcomes are likely due to differences in study design including the lengths of exposure or abstinence periods. Together, these findings show that acute and chronic ethanol exposure disrupts Hipp-NAc plasticity, providing a potential substrate for alcohol-induced alterations in reward-related behaviors.

## Acknowledgements

We would like to thank Dr. Yaohui Zhu and Kaela Befano for their assistance and Dr. Brian Mathur for his technical support, use of his Analox, and thoughtful comments on the manuscript. This work was supported by NSF IOS2402645, T32GM144876-02, the Meyerhoff program at UMBC, and start-up funds from UMBC.

## Notes

### Competing Interest Statement

The authors have declared no competing interest.

### Summary of Updates

Manuscript was edited. BEC data and an author were added.

## References

1. Koob, G. F. & Volkow, N. D. Neurocircuitry of Addiction. Neuropsychopharmacology 35, 217–238 (2010).

2. Lovinger, D. M. & Abrahao, K. P. Synaptic plasticity mechanisms common to learning and alcohol use disorder. Learn. Mem. Cold Spring Harb. N 25, 425–434 (2018).

3. Destoop, M., Morrens, M., Coppens, V. & Dom, G. Addiction, Anhedonia, and Comorbid Mood Disorder. A Narrative Review. Front. Psychiatry 10, 311 (2019).

4. Garfield, J. B. B., Lubman, D. I. & Yücel, M. Anhedonia in substance use disorders: A systematic review of its nature, course and clinical correlates. Aust. N. Z. J. Psychiatry 48, 36–51 (2014).

5. Russo, S. J. & Nestler, E. J. The brain reward circuitry in mood disorders. Nat. Rev. Neurosci. 14, 609–625 (2013).

6. Stuber, G. D., Britt, J. P. & Bonci, A. Optogenetic modulation of neural circuits that underlie reward seeking. Biol. Psychiatry 71, 1061–1067 (2012).

7. Carlezon, W. A. & Thomas, M. J. Biological substrates of reward and aversion: A nucleus accumbens activity hypothesis. Neuropharmacology 56, 122–132 (2009).

8. Calhoon, G. G. & O’Donnell, P. Closing the Gate in the Limbic Striatum: Prefrontal Suppression of Hippocampal and Thalamic Inputs. Neuron 78, 181–190 (2013).

9. Goto, Y. & Grace, A. A. Limbic and cortical information processing in the nucleus accumbens. Trends Neurosci. 31, 552–558 (2008).

10. O’Donnell, P. & Grace, A. A. Synaptic interactions among excitatory afferents to nucleus accumbens neurons: Hippocampal gating of prefrontal cortical input. J. Neurosci. 15, 3622–3639 (1995).

11. Volkow, N. D. & Morales, M. The Brain on Drugs: From Reward to Addiction. Cell 162, 712–725 (2015).

12. Scofield, M. D. et al. The nucleus accumbens: Mechanisms of addiction across drug classes reflect the importance of glutamate homeostasis. Pharmacol. Rev. 68, 816–871 (2016).

13. Francis, T. C. & Lobo, M. K. Emerging Role for Nucleus Accumbens Medium Spiny Neuron Subtypes in Depression. Biol. Psychiatry 81, 645–653 (2017).

14. Kircher, D. M., Aziz, H. C., Mangieri, R. A. & Morrisett, R. A. Ethanol Experience Enhances Glutamatergic Ventral Hippocampal Inputs to D1 Receptor-Expressing Medium Spiny Neurons in the Nucleus Accumbens Shell. J. Neurosci. 39, 2459–2469 (2019).

15. Renteria, R., Maier, E. Y., Buske, T. R. & Morrisett, R. A. Selective alterations of NMDAR function and plasticity in D1 and D2 medium spiny neurons in the nucleus accumbens shell following chronic intermittent ethanol exposure. Neuropharmacology 112, 164–171 (2017).

16. Lim, B. K., Huang, K. W., Grueter, B. A., Rothwell, P. E. & Malenka, R. C. Anhedonia requires MC4R-mediated synaptic adaptations in nucleus accumbens. Nature 487, 183–189 (2012).

17. Wilson, C. J. & Kawaguchi, Y. The origins of two-state spontaneous membrane potential fluctuations of neostriatal spiny neurons. J. Neurosci. 16, 2397–2410 (1996).

18. Stern, E. A., Jaeger, D. & Wilson, C. J. Membrane potential synchrony of simultaneously recorded striatal spiny neuronsin vivo. Nature 394, 475–478 (1998).

19. Goto, Y. & O’Donnell, P. Network synchrony in the nucleus accumbens in vivo. J. Neurosci. Off. J. Soc. Neurosci. 21, 4498–4504 (2001).

20. Floresco, S. B. & Phillips, A. G. Dopamine and hippocampal input to the nucleus accumbens play an essential role in the search for food in an unpredictable environment. Psychobiology 27, 277–286 (1999).

21. Ito, R., Robbins, T. W., Pennartz, C. M. & Everitt, B. J. Functional interaction between the hippocampus and nucleus accumbens shell is necessary for the acquisition of appetitive spatial context conditioning. J. Neurosci. 28, 6950–6959 (2008).

22. Britt, J. P. et al. Synaptic and Behavioral Profile of Multiple Glutamatergic Inputs to the Nucleus Accumbens. Neuron 76, 790–803 (2012).

23. Davidow, J. Y., Foerde, K., Galván, A. & Shohamy, D. An Upside to Reward Sensitivity: The Hippocampus Supports Enhanced Reinforcement Learning in Adolescence. Neuron 92, 93–99 (2016).

24. LeGates, T. A. et al. Reward behaviour is regulated by the strength of hippocampus–nucleus accumbens synapses. Nature 564, 258–262 (2018).

25. Sjulson, L., Peyrache, A., Cumpelik, A., Cassataro, D. & Buzsáki, G. Cocaine Place Conditioning Strengthens Location-Specific Hippocampal Coupling to the Nucleus Accumbens. Neuron 98, 926-934.e5 (2018).

26. Trouche, S. et al. A Hippocampus-Accumbens Tripartite Neuronal Motif Guides Appetitive Memory in Space. Cell 176, 1393-1406.e16 (2019).

27. Zhou, Y. et al. A ventral CA1 to nucleus accumbens core engram circuit mediates conditioned place preference for cocaine. Nat. Neurosci. 22, 1986–1999 (2019).

28. Heller, A. S. et al. Association between real-world experiential diversity and positive affect relates to hippocampal-striatal functional connectivity. Nat. Neurosci. 23, 800–804 (2020).

29. Huntley, E. D. et al. Adolescent substance use and functional connectivity between the ventral striatum and hippocampus. Behav. Brain Res. 390, 112678 (2020).

30. Sosa, M., Joo, H. R. & Frank, L. M. Dorsal and Ventral Hippocampal Sharp-Wave Ripples Activate Distinct Nucleus Accumbens Networks. Neuron 105, 725-741.e8 (2020).

31. Barnstedt, O., Mocellin, P. & Remy, S. A hippocampus-accumbens code guides goal-directed appetitive behavior. Nat. Commun. 15, 3196 (2024).

32. Griffin, W. C., Lopez, M. F., Woodward, J. J. & Becker, H. C. Alcohol dependence and the ventral hippocampal influence on alcohol drinking in male mice. Alcohol 106, 44–54 (2023).

33. Keith, L. D. & Crabbe, J. C. Specific and nonspecific effects of ethanol vapor on plasma corticosterone in mice. Alcohol 9, 529–533 (1992).

34. Eisenhardt, M., Hansson, A. C., Spanagel, R. & Bilbao, A. Chronic intermittent ethanol exposure in mice leads to an up-regulation of CRH/CRHR1 signaling. Alcohol. Clin. Exp. Res. 39, 752–762 (2015).

35. Chu, K., Koob, G. F., Cole, M., Zorrilla, E. P. & Roberts, A. J. Dependence-induced increases in ethanol self-administration in mice are blocked by the CRF1 receptor antagonist antalarmin and by CRF1 receptor knockout. Pharmacol. Biochem. Behav. 86, 813–821 (2007).

36. Lovinger, D. M., White, G. & Weight, F. F. NMDA receptor-mediated synaptic excitation selectively inhibited by ethanol in hippocampal slice from adult rat. J. Neurosci. 10, 1372–1379 (1990).

37. Hendricson, A. W., Thomas, M. P., Lippmann, M. J. & Morrisett, R. A. Suppression of L-type voltage-gated calcium channel-dependent synaptic plasticity by ethanol: analysis of miniature synaptic currents and dendritic calcium transients. J. Pharmacol. Exp. Ther. 307, 550–558 (2003).

38. Copenhaver, A. E. & LeGates, T. A. Sex-Specific Mechanisms Underlie Long-Term Potentiation at Hippocampus→Medium Spiny Neuron Synapses in the Medial Shell of the Nucleus Accumbens. J. Neurosci. 44, (2024).

39. Gill, K. M. & Grace, A. A. Differential effects of acute and repeated stress on hippocampus and amygdala inputs to the nucleus accumbens shell. Int. J. Neuropsychopharmacol. 16, 2013–2025 (2013).

40. Segev, A. & Akirav, I. Cannabinoids and Glucocorticoids in the Basolateral Amygdala Modulate Hippocampal–Accumbens Plasticity After Stress. Neuropsychopharmacology 41, 1066–1079 (2016).

41. Bryant, K. G. et al. A History of Low-Dose Ethanol Shifts the Role of Ventral Hippocampus during Reward Seeking in Male Mice. eNeuro 10, (2023).

42. Curran-Alfaro, C. M., Bryant, K. G., Ledgister, T.-S., Amin, S. & Barker, J. M. Sex differences in low-dose ethanol effects on motivated behavior and limbic corticostriatal activity in mice. Alcohol Clin. Exp. Res. 50, e70264 (2026).

43. Zorumski, C. F., Mennerick, S. & Izumi, Y. Acute and Chronic Effects of Ethanol on Learning-Related Synaptic Plasticity. Alcohol Fayettev. N 48, 1–17 (2014).

44. Ewin, S. E. et al. Chronic Intermittent Ethanol Exposure Selectively Increases Synaptic Excitability in the Ventral Domain of the Rat Hippocampus. Neuroscience 398, 144–157 (2019).

45. Van Skike, C. E., Goodlett, C. & Matthews, D. B. Acute alcohol and cognition: Remembering what it causes us to forget. Alcohol 79, 105–125 (2019).

46. Bach, E. C. et al. Chronic Ethanol Exposures Leads to a Negative Affective State in Female Rats That Is Accompanied by a Paradoxical Decrease in Ventral Hippocampus Excitability. Front. Neurosci. 15, (2021).

47. Izumi, Y., Nagashima, K., Murayama, K. & Zorumski, C. F. Acute effects of ethanol on hippocampal long-term potentiation and long-term depression are mediated by different mechanisms. Neuroscience 136, 509–517 (2005).

48. Jeanes, Z. M., Buske, T. R. & Morrisett, R. A. Cell type-specific synaptic encoding of ethanol exposure in the nucleus accumbens shell. Neuroscience 277, 184–195 (2014).

